# Genotype Imputation Performance of Three Reference Panels Using African Ancestry Individuals

**DOI:** 10.1101/245035

**Authors:** Candelaria Vergara, Margaret M. Parker, Liliana Franco, Michael H. Cho, Ana V. Valencia-Duarte, Terri H. Beaty, Priya Duggal

## Abstract

Genotype imputation is used to estimate unobserved genotypes from genome-wide maker data, to increase genome coverage and power for genome-wide association studies. Imputation has been most successful for European ancestry populations in which very large reference panels are available. Smaller subsets of African descent populations are available in 1000 Genomes (1000G), the Consortium on Asthma among African-Ancestry Populations in the Americas (CAAPA) and the Haplotype Reference Consortium (HRC). We aimed to compare the performance of these reference panels when imputing variation in 3,747 African Americans (AA) from 2 cohorts (HCV and COPDGene) genotyped using the Illumina Omni family of microarrays. The haplotypes of 2,504 individuals (from 1000G), 883 (from CAAPA) and 32,611 (from HRC) were used as reference. We compared the performance of these panels based on number of variants, imputation quality, imputation accuracy and coverage. In both cohorts, 1000G imputed 1.5–1.6x more variants compared to CAAPA and 1.2x more variants than HRC. Similar findings were observed for variants with higher imputation quality (R^2^>0.5) and for rare, low frequency, and common variants. When merging the results of the three panels the total number of imputed variants was 62M-63M with 20M overlapping variants imputed by all three panels, and a range of 5 to 15M unique variants imputed exclusively with one of the three panels. For overlapping variants, imputation quality was highest for HRC, followed by 1000G, then CAAPA, and improved as the minor allele frequency increased. The 1000G, HRC and CAAPA participants of African ancestry provided high performance and accuracy for imputation of African American admixed individuals, increasing the total number of variants with high quality available for subsequent analyses. These three panels are complementary and would benefit from the development of an integrated African reference panel, including data from multiple sources and populations.

## Introduction

Over the past 10 years, genome-wide association studies (GWAS) have uncovered a large number of replicated associations for many complex human diseases [1–3]. These studies have used different genotyping arrays with 300,000 to 2.5 million single nucleotide polymorphisms (SNPs), varied genomic coverage, and a wide range of allelic frequencies across populations. In general, these arrays provide excellent genomic coverage and density for European ancestry populations. Despite efforts at enrichment, imputation remains modest at best for other ancestral populations, especially populations of African ancestry. Genotype imputation is a cost-effective method for statistically predicting un-typed genotypes not directly assayed in a sample of individuals based on a dense reference panel of haplotypes. Imputation methods estimate haplotypes of observed genotypes shared between genotyped individuals and a sequenced reference panel, and use this information to infer alleles at un-typed SNPs [1]. This process can increase the overall genome coverage of an array by increasing the number of testable single nucleotide variants (SNVs) across the entire genome and can improve fine-mapping of a targeted region of interest. Imputation also facilitates the comparison and meta-analyses of studies originally done on different microarrays [1,2,4,5], potentially bridging the gap in coverage between various genome-wide SNP platforms [6].

Imputation of rare SNVs is more challenging since rare alleles are often ethnicity or population-specific and reflect fine-scale linkage disequilibrium (LD) structure impacted by recent demographic events [7]. Options for imputing low-frequency and rare variants more accurately in any specific population include increasing the size of the imputation reference panel to capture more reference haplotypes, or increasing the sequencing depth in the reference samples to minimize error rates inherent in low-coverage sequencing [8]. Recently admixed populations, which have higher degrees of LD and greater heterogeneity in their haplotype block structure (reflect the dynamics of admixture), may also benefit from using more diverse or larger reference populations.

Earlier available reference panels include the Human Genome Diversity Project [9], the HapMap Consortium [10] and the 1000 Genomes Project (1000G) [11]. More recently, the Haplotype Reference Consortium (HRC) [12] was constructed via a predominantly European ancestry consortium currently comprised of 32,611 individuals with whole genome or exome sequences available. The HRC includes the Genome of The Netherlands (GoNL), 250 Dutch parent-offspring families sequenced at 12x depth [13], the UK10K project with nearly 10,000 individuals whose whole genome was sequenced at 7x, or exome sequenced at 80x [14] and 1000G subjects among other cohorts (http://www.haplotype-referenceconsortium.org/participating-cohorts). Another project, funded by the UK government, plans to sequence 100,000 whole genomes from patients registered and treated by the National Health Service (http://www.genomicsengland.co.uk/the-100000-genomes-project/). These dense reference panels will allow better imputation of low frequency and rare variants [15] and the discovery of new variants [14,16], but are generally focused on populations of European descent.

There are a only few reference panels available for imputation in African Americans, those include the 1000 Genomes Project (1000G) [11] and the Consortium on Asthma among African ancestry Populations in the Americas (CAAPA) [17]. The 1000G includes 661 individuals with African ancestry from Esan, Gambian, Luhya, Mende, Yoruba, Barbadian and African-American populations [11]. The CAAPA panel is an additional resource completed on populations of African ancestry from the Americas [17]. CAAPA included 883 unrelated individuals of African descent from 15 locations in North, Central, and South America, the Caribbean, and Yoruba-speaking individuals from Nigeria. Their relatively small size of these panels compared to the references populations for European ancestry, limits the ability to discover new variants beyond those already present on the commercially available chips for subjects of African descent. Other projects assessing genetic diversity through dense genotyping and at the WGS level in African populations are currently under development including African Genome Variation Project (AGVP) [18] and the African Genome Resources (AGR) reference panel (https://www.apcdr.org/).

In this paper, we compared imputation performance using publicly available reference panels to evaluate imputation accuracy and quality in African or admixed populations of African descent. Imputation performance has been evaluated for African American populations comparing the 1000G, HapMap and the Exome Sequencing Project [5,19–22], and also using several combinations of populations from 1000G. Previous analyses suggest multi-ethnic panels in 1000G (primarily European (EUR) and African (AFR)) improve imputation performance compared to a reference panel from any single population (AFR) [21]. In previous studies of African Americans imputed with several combinations of 1000G populations, imputation accuracy (based on concordance and imputation quality score) was comparable across the reference panels. Imputation quality for SNPs with MAF between 0.02–0.50 was better when using more distantly related reference panels containing several continental African populations (AFR+EUR or ALL populations) in comparison with more closely related populations (Yoruba (YRI), CEPH European (CEU), and African Americans from the Southwest US (ASW)), but when analyzing all ranges of MAF including those with MAF < 0.02, the most closely related (YRI+CEU+ASW) panel produced better imputation results. On the other hand, genotype concordance was similar for both distant and closely related reference panels from 1000G [5].

Imputation is standard part of all array-based genome-wide association analyses. However, the relative performance of these newer imputation reference panels – with varying total sample size and number of African individuals – is unknown. In this study, we extend these prior imputation comparisons beyond the 1000G populations by evaluating genotype imputation performance using CAAPA, HRC, and 1000G reference panels in two independent populations of African Americans [23–25].

## Methods

The current study includes a total of 3,747 African Americans participating in previous genome-wide association studies of spontaneous resolution of Hepatitis C viral infection (HCV cohort) and Chronic Obstructive Pulmonary Disease (COPD) from the COPDGene cohort, a multi-site study of heavy smokers [23–25]. Metrics of imputation performance, accuracy, genome coverage and annotation of variants were calculated in these two cohorts, separately.

### Study subjects

<u>African Americans from the HCV cohort:</u> A genome-wide marker panel from 447 African Americans was used, as previously described [23,24]. Briefly, 2,401 African American individuals participating in a longitudinal cohort study or identified through blood repositories as having HCV infection (spontaneously resolved or persistent) were enrolled, and were genotyped as part of the HCV Genetics Consortium. Each individual study obtained consent for genetic testing from their governing Institutional Review Board (IRB) and the Johns Hopkins School of Medicine Institutional Review Board.

<u>African Americans from the COPDGene cohort.</u> This study included 3,300 African Americans participants in the COPDGene study. A complete study protocol for COPDGene had been described elsewhere [25]. Briefly, 10,280 self-identified Non-Hispanic Whites and African Americans between the ages of 45 and 80 years with a minimum of 10 pack-years smoking history were enrolled at 21 centers across the US with the goal of identifying genetic causes of COPD. Each study site has obtained local IRB approval to enroll participants in this project, and all subjects provide informed consent [25].

### Genotyping and Quality Control

<u>African Americans from the HCV cohort:</u> Genetic variants and their locations for the genotypic data, reference panels and whole genome sequencing data were specified based on The Genome Reference Consortium Human build 37 (GRCh37) [26]. A total of 774,792 SNPs genotyped on the Illumina Omni Quad array (IIlumina, Inc. San Diego) met quality control criteria and were used for imputation. SNPs with MAF < 0.01, those with missing call rate ≥ 5% and those deviating from Hardy Weinberg equilibrium at p< 1×10^−5^ were removed from the analysis for quality control. Individuals cryptically related, duplicated replicates, and individuals with sex discrepancies were excluded [23].

<u>African Americans from the COPDGene cohort:</u> A total of 624,564 SNPs genotyped on the Illumina Omni Express array (IIlumina Inc. San Diego, CA) were used for imputation. All SNPs with MAF < 0.05, those with missing call rate ≥ 2%, those deviating from Hardy Weinberg equilibrium at p< 1×10^−3^ and individuals cryptically related, duplicated replicates and individuals with sex discrepancies were excluded [27].

### Whole-Genome Sequencing: Library Preparation and Bioinformatic Analysis

Whole genome sequencing data in a subgroup of 17 subjects of the HCV cohort was performed at the New York Genome Center. In brief, libraries of 350-bp fragments were generated from 1 µg sheared genomic DNA using the TruSeq PCR-Free library preparation kit (Illumina, San Diego, CA). WGS was performed at a coverage of 30x. Base calling and filtering were performed using current Illumina software; sequences were aligned to NCBI genome (build 37) using Burrows-Wheeler Aligner [28]; Picard was used to remove duplicate reads [29]; base quality scores were recalibrated using GATK [30]. Assessment of reads not aligning fully to the reference genome was performed, locally realigning around indels to identify putative insertions or deletions in the region. Variants were called using GATK HaplotypeCaller tool, which generates single-sample Genomic VCF (GVCF) files. To improve variant call accuracy, multiple single-sample GVCF files were jointly genotyped using GATK Genotype GVCFs, which generates a multi-sample VCF. Variant Quality Score Recalibration (VQSR) was performed on the multi-sample VCF, which adds quality metrics to each variant that can be used in downstream variant filtering [31]. Quality control of all variants included filtering out based on genotyping quality score <20, read depth <10 and removing variants in genomic duplicated segments.

### Estimation of Genetic African American Ancestry

Genetic ancestry for both cohorts was determined by principal components using the *smartpca* program in EIGENSOFT [32]. A subset of independent SNPs across the genome were selected by pruning the full dataset for markers with an *r^2^* < 0.01 to insure independence between SNPs. Chromosomal regions known to be associated with ethnicity were removed (including the lactase regions on chromosomes 2, 8, and the HLA region on chromosome 6). African-American ancestry groups were determined based on tight clustering of the first 2 principal components. Outliers were removed based on heterozygosity, and if the subjects were 6 standard deviations from either of the 2 first principal components [23,25] The average percent of African ancestry was estimated at 79.5% and 80.1% in the HCV and COPDGene cohorts, respectively.

### Phasing and Imputation

For both cohorts, Eagle v2.3, a reference-based phasing algorithm was used to phase genotypes prior to imputation [33,34]. Imputation was performed for chromosomes 1 to 22 using the Minimac3 software through the publicly available Michigan Imputation Server [4]. Minimac3 is a Markov Chain based haplotyper that can resolve long haplotypes or infer missing genotypes in samples of unrelated individuals [35]. Imputation of genotypes was performed using 3 different reference panels:

a) **1000 Genomes Phase 3, Version 5** (referred to here as “1000G”) included 49,143,605 sites located in chromosomes 1 to 22 for the complete set of 2,504 individuals representing 5 continental and sub-continental populations: East Asian (EAS= 504), European (EUR=503), South Asian (SAS=489), African (AFR= 661) and Mixed American (AMR= 347) [11]. The 1000G uses a combination of low-coverage whole-genome sequencing (WGS) (mean depth of 7.4x), deep exome sequencing (mean depth of 65.7x) and dense microarray genotyping;
b) **The CAAPA reference panel** (“CAAPA”) comprising 883 individuals from 19 case-control studies of asthma with 31,163,897 variants identified on chromosomes 1–22 by high coverage WGS (30x). The populations for this panel include individuals from Barbados (N=39), Jamaica (N=45), Dominican Republic (N=47), Honduras (N=41), Colombia (N=31), Puerto Rico (N= 53), Brazil (N=33) and Nigeria (N=25) and African Americans (N=328) [17];
c) **The Haplotype Reference Consortium** (“HRC”) reference panel combining data sets from 20 different studies with low-coverage WGS (4–8x coverage) of subjects with predominantly European ancestry. The first release of this reference panel consists of 32,611 individuals with 64,976 haplotypes including 39,235,157 SNPs, each with a minor allele count (MAC) greater or equal to 5 [12].

### Imputation Performance Metrics

<u>Evaluation of imputed variants by reference panel:</u> For each reference panel and cohort, we assessed imputation performance using the following criteria: 1) the total number of imputed variants; 2) the distribution of all variants based on MAF ranges; and 3) the relationship between imputation quality and MAF. Imputation quality was determined using the R^2^ score, or the estimated value of the squared correlation between imputed genotypes and true, unobserved genotypes basing its calculation in the population allelic frequencies [36].

<u>Comparison of imputed variants between reference panels:</u> To compare imputation results between panels, we analyzed variants imputed by all three panels (“overlapping” variants) and exclusively by each panel (“unique” variants). For overlapping variants, the imputation quality and genotype concordance between the panels was compared. Unique variants with R^2^ >0.5, were evaluated by their number and MAF and for its presence and MAF in the CEU, YRI and CHB populations from1000G as a method to evaluate the potential ancestral origin of them.

### Imputation accuracy

Imputation accuracy is defined as the proportion of correctly imputed SNPs among all successfully imputed SNPs [37–40]. We calculated imputation accuracy using three separate approaches:

a) A “masked analysis” where we removed genotypes of a subset of 25,000 SNPs and then imputed them as though they were not genotyped. Imputed genotypes of these SNPs then were compared back to their original genotypes [5,37,41]. The MAF of these SNPs ranged from 0.01 to 0.5;
b) A comparison of the allelic dosage of the original genotypic data with the allelic dosage of imputed data. Given three genotypes AA, AB, and BB for each SNP, the allelic dosage for each individual can be calculated as probabilities (P) of each of three genotypes via 2*P (AA) + 1*P(AB) + 0*P(BB) to obtain the expected allelic dosage from the original genotypic data and from the observed allelic dosage for masked and imputed genotype at each SNP [42]. The metric EmpRsq obtained in Minimac3 is the correlation between the true genotyped values and the imputed dosages calculated by hiding all known genotypes for a given SNP [36], similar to the masked analysis described above. We calculated the mean of this EmpRsq by bins of 0.001–0.01 value of frequency of the minor allele.
c) A comparison of the imputed genotypes with whole genome sequencing genotypes in a subgroup of 17 individuals from the HCV study. This analysis was restricted to variants located on chromosome 22, and was done independently for all the variants imputed with each reference panel.

### Genomic Coverage and Density of Imputed Variants

The total proportion of genomic variation captured by an array, either directly or indirectly, is referred to as “genomic coverage.” Assessments of imputation-based genomic coverage leverages observed array SNPs which imputed from a more densely genotyped or sequenced reference panel, such as the HapMap Project3 or 1000 Genomes Project [43,44]. In this study we based our calculations on the imputation R^2^ (calculated as squared correlation between the actual (discrete) allelic dosage at a SNP and the imputed (continuous) allelic dosage, over a defined set of samples). Genomic coverage was quantified as the proportion of variants with an imputation R^2^ ≥ 0.8, and the reference set of variants used to determine imputation-based genomic coverage was the total number of variants described in each imputation reference panel. This method has been described and used previously as one assessment of genomic coverage in imputation performance studies [45,46].

We also calculated the density of imputed variants (represented as the number of variants with R^2^>0.8 per Kb) across all autosomes, by chromosomes and in chromosomal regions harboring known genes. We compared the results obtained with the three panels. Variants genotyped on the arrays were given imputation R^2^ = 1 for all coverage and density calculations; chromosome sizes in base pairs were obtained from the UCSC Known Gene Human Annotation (GRCh37). Coordinates of the regions containing genes were obtained from the RefSeq database via UCSC genome browser (http://genome.ucsc.edu/). Plink version 1.90 beta [47], bcftools [48] and customized scripts in R [49] were used for the analyses with both cohorts.

## Results

### Imputation Performance Metrics

<u>Evaluation of imputed variants by reference panel:</u> The total number of imputed variants and their distribution by MAF was similar for both the HCV and COPDGene cohorts (Tables 1 and 2). In the HCV cohort, 1000G imputed 1.5x more variants than did CAAPA, and 1.2x more variants than did HRC (46,626,711 vs. 29,850,712 and 39,019,013, respectively) regardless of imputation quality. This difference was similar for the COPDGene cohort at 1.6x and 1.2x, respectively. However, it is important to note the 1000G imputation includes small insertions/deletions (INDELS) that are not currently available in the HRC and CAAPA panels. These INDELS correspond to 3,109,956 variants for the HCV cohort (6.7% of total) and 3,303,782 for the COPDGene cohort (7.0 % of total).

**Table 1.** Number of variants imputed by reference panel, minor allele frequency ranges and imputation quality for the HCV cohort.

|  | CAAPA |  | 1000G |  | HRC |  |
| --- | --- | --- | --- | --- | --- | --- |
| Minor allele frequency | Number of variants | $R^2 > 0.5$ (%) | Number of variants | $R^2 > 0.5$ (%) | Number of variants | $R^2 > 0.5$ (%) |
| 0-0.0001 | 12,164 | 0 | 396,072 | 0 | 3,568,020 | 0 |
| 0.0001-0.01 | 15,481,830 | 36.8 | 29,683,849 | 33.9 | 21,736,936 | 32.1 |
| 0.01-0.05 | 6,343,792 | 83.9 | 7,050,809 | 97.4 | 6,055,818 | 99.3 |
| >0.05 | 8,012,926 | 94.2 | 9,495,981 | 98.6 | 7,658,238 | 99.7 |
| Total | 29,850,712 | 62.3 | 46,626,711 | 56.4 | 39,019,012 | 52.9 |

**Table 2.** Number of variants imputed by reference panel, minor allele frequency ranges and imputation quality for the COPDGene Cohort.

|  | CAAPA |  | 1000G |  | HRC |  |
| --- | --- | --- | --- | --- | --- | --- |
| Minor allele frequency | Number of variants | R<sup>2</sup> > 0.5 (%) | Number of variants | R<sup>2</sup> > 0.5 (%) | Number of variants | R<sup>2</sup> > 0.5 (%) |
| 0-0.0001 | 374 | 0 | 60,632 | 0 | 1,119,183 | 0 |
| 0.0001-0.01 | 15,528,412 | 38.4 | 30,331,067 | 41.52 | 24,294,755 | 41.1 |
| 0.01-0.05 | 6,309,636 | 81.02 | 7,085,542 | 96.94 | 6,038,380 | 99.1 |
| >0.05 | 8,007,965 | 93.6 | 9,624,269 | 98.4 | 7,675,728 | 99.7 |
| Total | 29,846,387 | 62.2 | 47,101,510 | 61.4 | 39,128,046 | 60.4 |

For variants imputed with R^2^ > 0.5, 1000G imputed 1.4x and 1.3x more variants (26,310,578 vs. 18,584,433 and 20,643,333 for CAAPA and HRC, respectively) in the HCV cohort. This same observation was seen in the COPDGene cohort where 1000G imputed 1.5x and 1.2x more variants than did either CAAPA or HRC. All three reference panels had a similar percent of variants imputed with R^2^ > 0.5 in both cohorts: HCV cohort (HRC, 53%; 1000G, 56%; CAAPA, 62%); COPDGene cohort (HRC, 60%, 1000G, 61%, CAAPA, 62%). For both cohorts and panels, the percentage of variants imputed with R^2^ > 0.5 increased with increasing MAF (Tables 1 and 2).

Regardless of allele frequencies, the number of imputed variants was greater in 1000G than for CAAPA: 1.8x more rare variants (MAF = 0.0001–0.01), 1.3x more low MAF variants (MAF=0.01–0.05) and 1.2x more common variants (MAF>0.05). These numbers were also higher for 1000G compared to HRC 1.4x, 1.1x and 1.2x, respectively. Similar values were observed in the COPDGene cohort, 1000G imputed 1.9x and 1.2x more rare variants than obtained with CAAPA and HRC, respectively. Those values were 1.1x more for low MAF variants and, 1.3x (1000G vs. CAAPA) and 1.2x (1000G vs. HRC) for the comparisons of the panels for common variants (Table 2). The distribution of the number of variants with R^2^ > 0.5 by MAF for each panel was similar between all reference panels with a high number of low frequency SNPs (i.e. those with MAF < 0.1) in the three panels for both cohorts.

For both cohorts and all three panels, imputation quality improved as the MAF increased, reaching a mean quality score or R^2^ of 0.6 or higher for common variants (MAF>0.05). CAAPA imputed with slightly lower quality across all MAF followed by 1000G and HRC (Figure 1). The higher imputation quality observed with HRC and 1000G was particularly evident in low frequency variants (i.e. those with MAF from 0.002 to 0.05). In the COPDGene cohort, HRC had a better performance compared to 1000G for very rare variants (MAF < 0.001) but this was not observed in the HCV cohort, likely due to sample size differences in the two target populations.

**Figure 1.**
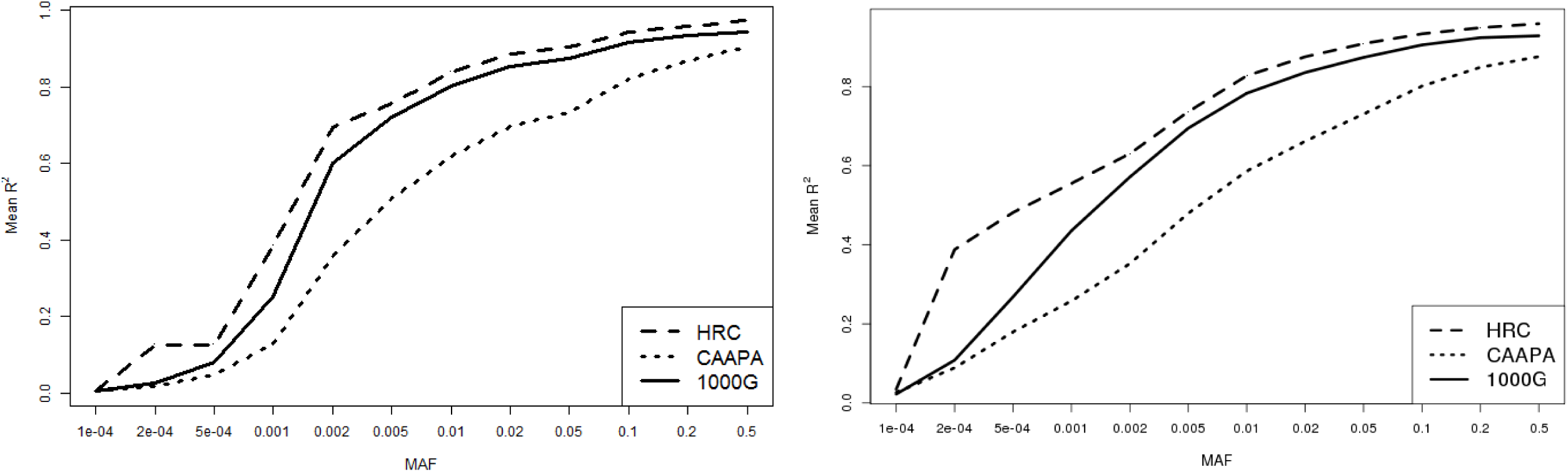
Relationship between imputation quality and minor allele frequency for all variants imputed with 1000G, CAAPA and HRC in the HCV cohort (left) and COPDGene cohort (right). The graph represent the mean of imputation R^2^ by intervals of minor allele frequency (0.001–0.01).

<u>Comparison of imputed variants between reference panels</u>: When merging the variants imputed independently with each reference panel, the total number of imputed variants was 62,673,539 for the HCV cohort and 63,525,310 for COPDGene cohort, representing an increase of 20 - 30 million variants compared to the imputation of each panel separately (Figure 2 and Supplementary Table 1). In both cohorts, there were approximately 20 million overlapping variants imputed with all three reference panels and a range of 5 to 15 million unique variants imputed exclusively within one of the three panels.

**Figure 2.**
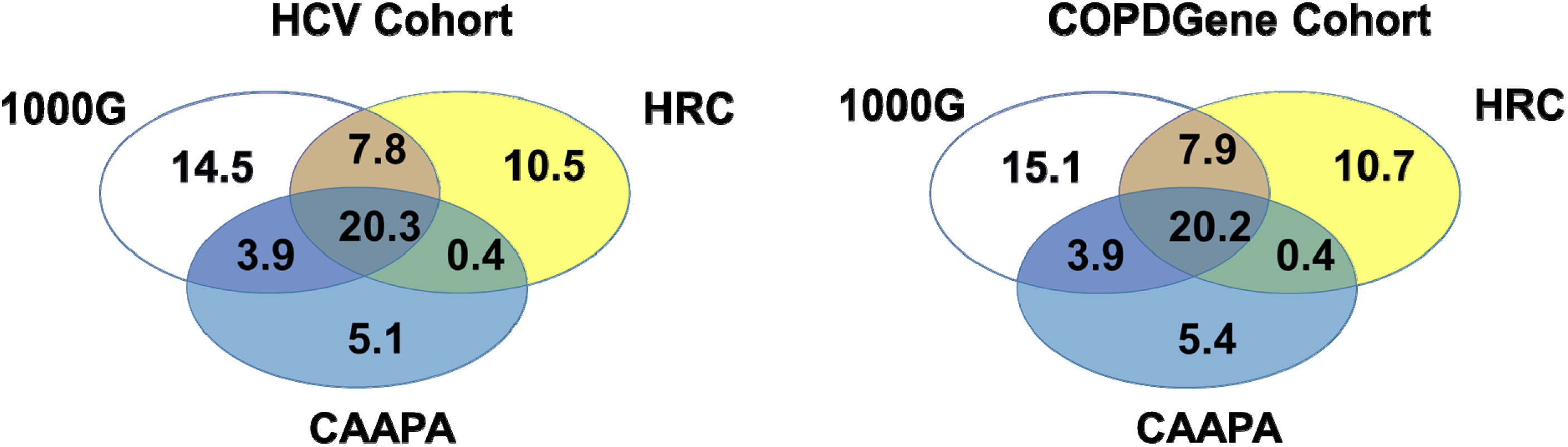
Number of overlapping and unique variants imputed with 1000G, CAAPA and HRC for the HCV cohor ort and COPDGene cohort. The intersection shows the number of variants imputed with the three reference panels and the non-overlapping sections of the circles show the variants unique to each panel.

For overlapping variants, we compared the imputation quality obtained with each of the three panels (Figure 3). For the same variants, the imputation quality was higher for HRC and 1000G compared to imputation run against CAAPA. From approximately 20 million overlapping variants, HRC and 1000G imputed ~18–19 million variants with R^2^ > 0.5, whereas CAAPA imputed ~15 million in both cohorts (Figure 2). Genotype calls of overlapping variants were 98–99% concordant between pairs of panels in both, COPDGene and HCV cohorts.

**Figure 3.**
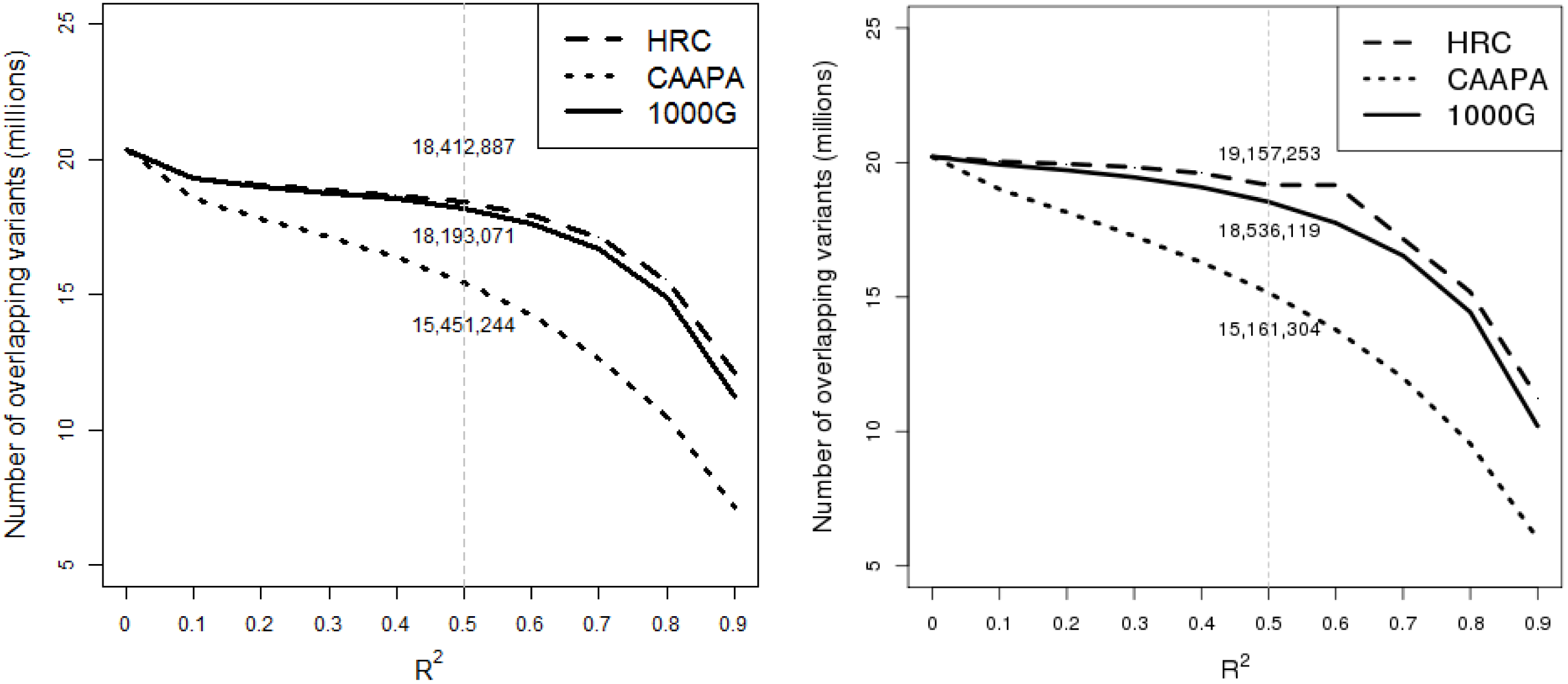
Comparison of imputation quality (R^2^) of all overlapping variants imputed with CAAPA, 1000G and HRC for the HCV cohort (left panel) and COPDGene cohort (right panel). The values on the gray line at imputation R^2^=0.5 correspond to the number of overlapping variants imputed with R^2^>= 0.5 with each panel.

In the HCV cohort, unique variants corresponded to 17.3%, 31.1% and 26.9% of all variants imputed with in CAAPA, 1000G and HRC, respectively. In the COPDGene cohort they were 17.9%, 32.1% and 27.3%, respectively. Most of them had MAF < 0.01 (75%-90%) in both cohorts for the three panels (Supplementary Tables 2 and 3), 5–24% of these rare variants were imputed with R^2^ > 0.5. There was a lower percentage (0.2–5%) of low frequency variants (MAF between 0.01–0.05) that were imputed with better quality (> 45% had R^2^> 0.5). The percentage of all unique variants imputed with R2 > 0.5 in the HCV cohorts was 27% for 1000G; 24% for CAAPA and 4% for HRC. A similar number of total unique imputed variants was observed in the COPDGene cohort: 34% for 1000G, 27% for CAAPA and 14% for HRC.

We interrogated the three parental populations of 1000G (CEU, YRI and CHB) to estimate allele frequencies of unique variants imputed with R^2^ > 0.5 in each population. 31% of the variants imputed with CAAPA were present in parental populations of 1000G. Only a small percentage of those were polymorphic in the CEU and CHB (0.3%-6% in both cohorts) as compared to YRI (12% for HCV and 22% for COPDGene - Supplementary Figure 1). Of those unique variants imputed using HRC, 34–35% were present in the parental populations; from those, 17–18% were polymorphic in CEU, in contrast with a 0.5–4% and 2% of variants that were polymorphic in YRI and CHB in both the HCV and COPDGene cohorts. All the unique variants with R^2^ > 0.5 imputed with 1000G in the HCV cohort were also present in the parental populations, and a large number were actually polymorphic (i.e. MAF > 0) in YRI (62%), CEU (36%), and CHB (31%). Similarly, in the COPDGene cohort, all unique variants imputed with 1000G were present in the three parental populations: YRI (56%), CEU (32%) and CHB (28%).

### Imputation accuracy

The concordance of genotype calls between the original genotype data and imputed data was quite high (96–97%) across all three panels using masked SNPs from both cohorts. The correlations between dosages of the imputed genotypes and actual genotypes ranged from 0.80–0.94, with higher correlations occurring when the MAF was greater than 1% (Supplementary Figure 2). In addition, we evaluated the concordance between whole genome sequenced and imputed data using variants on chromosome 22 in the HCV cohort. The genotype concordance for the three panels was 99% for 213,467, 190,005, and 195,591 variants overlapping between sequenced genotypes and imputed genotypes with 1000G, CAAPA and HRC, respectively.

### Genomic Coverage and Density of Imputed Variants

We imputed 20,222,182 variants with R^2^> 0.8 using the 1000G panel, 11,684,700 with CAAPA and 16,941,215 with HRC in the HCV cohort. In the COPDgene cohort, we obtained 19,645,517; 10,558,297 and 16,831,447 variants with those three panels, respectively. The genomic coverage was 0.41, 0.37 and 0.43 for 1000G, CAAPA and HRC, respectively in the HCV cohort and 0.39, 0.33 and 0.42 in the COPDGene Cohort. Genomic density of markers included in the genotype array was estimated to be at 0.3 and 0.2 marker/Kb in HCV cohort and COPDGene cohort respectively. In contrast, imputation with 1000G, CAAPA and HRC increased the genome density to 7, 4.1 and 5.9 markers per Kb, respectively. For both cohorts, the average density across chromosomes was ~6.8 variants/Kb for 1000G, ~3.9 variants/Kb for CAAPA and ~5.7 variants/Kb for HRC. The density was considerably lower in gene regions with an average ~1.4, ~2.3 and ~1.9 variants/Kb for CAAPA, 1000G and HRC, respectively (Supplementary Table 3 and 4).

### Discussion

In the current study, we used three reference panels to impute GWAS genotyping data in two independent cohorts of African American individuals with remarkably consistent results between the two studies. Imputation to three reference panels increased coverage and density of markers across all autosomal chromosomes and facilitated the accurate imputation of both rare and common alleles with R^2^ > 0.5. Somewhat surprisingly, despite the smaller size of the reference panel and number of African-Americans, the 1000G reference panel resulted in a higher number of imputed variants (even after removing INDELS) than either the HRC or CAAPA cohorts alone. Additionally, while all three panels led to accurate estimated genotypes, the imputation quality was highest for HRC across all MAFs, but especially for low frequency and rare variants.

A greater number of variants were imputed with 1000G as compared to CAAPA. The substantially larger sample size of the 1000G panel may explain this difference by itself. Previous studies comparing references panels have shown larger reference panels considerably increase the number of imputed variants, as well as their imputation quality and accuracy, particularly for low-frequency variants [8,12,15]. However, the composition of the reference population and similarity to the target population is also very important. Shriner et al. [41] imputed variants on chromosome 22 in African Americans from the Washington, D.C. metropolitan area participating in the Howard University Family Study [50] and concluded the YRI reference panel outperformed other HapMap reference panels, including ASW, as well as a combination of the CEU and YRI. Previous studies in European populations have indicated population specific panels can improve imputation quality and coverage [51,52] compared to broader panels. This improvement in the number of variants imputed and the accuracy using population specific panels argues that LD patterns of ethnic-specific variants may not be captured by different ethnic groups with distinct ancestral genetic background [53], which might include haplotypes from irrelevant populations. We would expect a population specific panel like CAAPA, where ~50% are African Americans with African ancestry estimates of 76% or higher [17], would be optimal for imputing more rare and common variants with higher quality in a target sample with similar high proportion of African ancestry (African ancestry estimates averaging 79.5%) [23,24]. However, in our study, the higher number of individuals from populations from continental Africa contained in 1000G compared to CAAPA (N=504 vs. N=25) may have outweighed the larger number of total African ancestry individuals in CAAPA, and provided more information on parental haplotype diversity [54,55] improving the chances of a rare variant being effectively tagged by a characteristic haplotype in admixed individuals [44].

1000G also imputed more variants in our African American subjects than did HRC even after accounting for indels, (even though 1000G is contained within this larger reference panel). The predominantly European ancestry haplotypes of the HRC panel might impair the selection of optimal haplotypes for imputation of these populations with high proportion of African ancestry and consequently explain why there was a less of an increase in imputation success in them. Our sensitivity analysis investigating the potential ancestral origin of the unique variants indicated 1000G was able to impute variants of European and African origin compared to the HRC panel alone where the unique variants were present mostly in the CEU population of 1000G indicating an exclusive European origin. This may also explain why HRC imputed more variants than obtained with the CAAPA reference panel alone, if the higher number of imputed variants obtained with HRC compared to CAAPA reflects underlying European haplotypes in admixed African Americans.

Regardless of reference panel, imputation yielded accurate genotypes as shown in the analysis of correlation between true genotypes and imputed genotypes, the masked analysis and the concordance analysis comparing imputed SNPs and sequencing data in the HCV cohort. The accuracy reflected in the estimated correlation of true genotypes and imputed genotypes was comparable (but slightly higher) for 1000G when studied as a function of minor allele frequency. The lower accuracy found using the CAAPA and HRC reference panels separately might be due to the inclusion of admixed populations without a large African and European ancestral panel in CAAPA or the predominance of European haplotypes in HRC. This could limit selection of the best reference haplotypes.

In this study we used global concordance as a measure of imputation accuracy excluding variants with MAF < 0.01. Accuracy can be inflated when calculations of concordance rate include rare and low frequency variants, due to chance concordance or chance agreement [56,57]. Due to the baseline low allele frequency, there is a low probability of any rare allele being present in any imputed sample; therefore, when the major allele is assigned in imputation, this inference would be almost uniformly “correct” by chance alone. This inflation is increasingly problematic whenever studies are more interested in evaluating low frequency and rare variants. Since our study didn’t include rare variants in the estimate, we consider our obtained values reliable. Our global estimates of accuracy were higher than previous results obtained in a group of 40 African Americans imputed with MACH using CEU, YRI, MEX and JPT+CHB HapMap populations. Accuracy values (measured as percentage of most likely genotypes agreeing with the original genotypes) of 88.8, 87.9 and 87.2 were found when masking 50%, 70% and 100% of all high quality SNPs [58]. The differences between this study and the current analysis are likely due to our larger sample size and type of reference population (Hapmap vs 1000G, HRC and CAAPA, separately).

Imputation resulted in several rare and common variants unique to CAAPA, although they are present in the 1000G database. These variants are predominantly of African origin even though a great number are monomorphic in YRI subjects. It is likely these unique variants may be derived from African genomes not included in the 1000G, and may be unique to African descent populations in the Americas (where there is also a small percentage of Native American genes included). Similarly, the unique variants imputed with HRC may correspond to European derived polymorphisms not captured by the haplotype structure of the other reference panels.

In this study, the total number of imputed variants increased when merging imputed variants obtained from each reference panel individually. However, although all panels are publicly available, we were not able to evaluate the imputation of a fully integrated reference panel. Previously integration of the 1000G and African Genome Variation Project panels markedly improved imputation accuracy across the entire allele frequency spectrum for populations poorly represented in the 1000G panel [18]. Similar results were found when merging the Estonian Biobank of the Estonian Genome Center, University of Tartu (EGCUT) and 1000G datasets [51] and the GoNL and 1000G [52] and when using a combined reference panel of 1,092 subjects from 1000G and 3,781 from UK10K Project for imputing rare variants in the Framingham Heart Study and the North Chinese Study [53]. We encourage development more publically available combined reference panels, like HRC, for African ancestry populations. This should also include the NHLBI funded Trans-Omics for Precision Medicine (TOPMed) project [59]. The 1000 Genomes Project will soon become “The International Genome Sample Resource” with all sequenced reads being re-mapped to the GRCh38 map producing new variants calls specific to this assembly. It will also expand the global catalogue of freely available sequence information by incorporating Russian samples, new African populations and whole genome sequences from the Simons Genome Diversity Project [60]. Data from the CAAPA project is also available at the database of Genotypes and Phenotypes (dbGaP) and can be used to explore the option of “custom reference panels” for imputation in African Americans and other admixed populations from Latin America and the Caribbean. But as we show in this study, there is still a need for characterizing large, diverse parental populations such as those from sub-Saharan African to better capture populations such as those in the Americas.

In summary, we found the 1000G, HRC and CAAPA reference panels provide high performance and accuracy for imputing dense marker panels for admixed African American individuals, increasing the total number of high quality imputed variants available for subsequent analyses. The 1000G panel also showed higher performance compared to the HRC and CAAPA reference populations likely because it included more diverse parental populations. Finally, there are a large number of variants unique to these three reference panels, making them complementary to each other. We recommend directing efforts to the construction of an integrated African panel including data from multiple resources and populations.

